# Subcellular Mass Spectrometry Reveals Proteome Remodeling in an Asymmetrically Dividing (Frog) Embryonic Stem Cell

**DOI:** 10.1101/2025.05.12.652541

**Authors:** Bowen Shen, Leena R. Pade, Fei Zhou, Peter Nemes

## Abstract

Subcellular proteomics holds the potential to reveal the molecular architecture of cellular processes with unprecedented spatial resolution. Performing these analyses deeply, at the level of hundreds to thousands of proteins in subcellular resolution, is a high and still unmet technical need. Here, we advance microprobe capillary electrophoresis–mass spectrometry (CE-MS) to achieve deep proteomic coverage—quantifying over 1,000 proteins within opposing poles of an asymmetrically dividing embryonic stem cell (blastomere). We integrated CE-electrospray ionization (CE-ESI) with trapped ion mobility spectrometry time-of-flight (timsTOF) MS, implementing data-independent acquisition (DIA) via parallel accumulation–serial fragmentation (diaPASEF). This CE-diaPASEF workflow identified 1,035 proteins from ∼200 pg of proteome digest, equivalent to ∼80% of the HeLa cell’s content, with high reproducibility (coefficient of variation <15% across technical triplicates). With microprobe sampling, this technology quantified 808 to 1,022 proteins in opposing poles of a dorsal-animal (D1) blastomere in the 8-cell *Xenopus laevis* embryo. Comparative proteomic analysis of the D1 blastomere and its descendants—the dorsal-animal-midline (D11) and dorsal-animal-lateral (D12) cells—revealed diverse molecular outcomes of asymmetric division: some protein profiles remained conserved, while others underwent significant or even reversed changes as these lineages descended into neural tissue and epidermal trajectories. Ultraviolet light-induced ventralization was performed to help disentangle subcellular gradients from dorsal-ventral patterning. Collectively, this work establishes microprobe CE-diaPASEF as a powerful platform for deep subcellular proteomics, enabling new insights into spatial proteome organization during key developmental processes.

## INTRODUCTION

Understanding the cell requires not only identifying its overall molecular components but also mapping their precise spatial organization and dynamic interactions. Subcellular molecular characterization is essential for decoding the processes governing cellular function in developing health and disease states. The proteome, the comprehensive repertoire of proteins expressed within a cell, offers a uniquely functional and high-resolution read on cellular activity. Yet, while imaging techniques such as super-resolution optical and cryo-electron microscopy (reviewed in [1]) revealed stunning spatial details, they are limited in throughput, typically detecting only a handful of proteins at a time. Mass spectrometry (MS)-based proteomics, on the other hand, can quantify thousands of proteins but has traditionally lacked spatial resolution. Bridging this gap— achieving deep proteomic profiling with subcellular resolution—represents a critical frontier in cell biology, enabling comprehensive insights into the spatial organization of cellular processes.

Single-cell mass spectrometry (MS) rapidly emerged to fill this analytical gap. The field’s current state was the focus of recent reviews (e.g., [2]). Following the inception of targeted hemoglobin detection in single erythrocytes[3] and carbonic anhydrase[4] in approximating amounts, the detection of ∼800 proteins marked the inception of the discovery of single-cell MS in dissected giant embryonic stem cells (blastomeres) in the South African clawed frog (*X. laevis*)[5]. These developments benefited from advances in capillary electrophoresis (CE)[6] and nano-flow liquid chromatography (nanoLC)^[5c,^ 7^]^, which improved sensitivity to ∼1,600 proteins, quantifying cell heterogeneity among identified embryonic cells and their lineages^[6b]^. Cell isolation, processing, and separation were downscaled to smaller cells, using automation recently to achieve high throughput (reviewed in ^[2c, 2f, 8]^). Modern nanoLC-MS can routinely report ∼3,000 proteins per somatic cell in broad biological contexts, e.g., recently including early metastatic lung cells[9] and monocytes/macrophages[10] to study cell heterogeneity. Patch-clamp proteomics[11] upscaled to the whole neuron, and analysis of the entire cell’s proteome improved sensitivity to ∼1,400 proteins using nanoLC-MS.[12] These studies provide ever-deepening coverage of single-cell proteomes at the level of the entire cell, some bordering subcellular proteomics.

Recent advances in subcellular MS are beginning to extend beyond small molecules and peptides (in part reviewed in [13]) into the realm of proteins. However, challenges in subcellular sampling and efficiency in molecular extraction and processing continue to limit sensitivity. Many current approaches successfully navigate the challenge via subcellular enrichment strategies on millions of cells, detecting thousands of proteins[14] but at the expense of losing single-cell or cell-type-specific context.[15] For example, large-scale MS combined with immunofluorescence microscopy enabled the subcellular localization of over 12,000 human proteins across 13 major organelles.[16] Similarly, nanoLC-MS, following immunoprecipitation, quantified more than 7,600 proteins from 19 compartments.[17] A high-throughput workflow that simultaneously tracks proteome and phosphoproteome dynamics across six distinct subcellular compartments found highly compartmentalized signaling responses to stimuli.[18] While these approaches deliver impressive depth and compartmental specificity, they remain dependent on large cell populations—underscoring the need for next-generation technologies capable of deep, spatially resolved proteomics within individual cells.

To meet this challenge, only recently have microsampling approaches emerged for high-sensitivity MS. We pioneered precision-fabricated microcapillaries to aspirate cell contents from 2-cell zebrafish (*Danio* rerio) and 16–128-cell *Xenopus laevis* embryos, identifying ∼800 proteins *in situ* ^[6b]^ and *in vivo*^[6a]^ (reviewed in part in ^[13b, 19]^). Patch-clamp proteomics further extended these capabilities to ∼35-µm-diameter neurons in live mouse brain slices, yielding ∼160 proteins from 1 pg, less than 0.25% of the total soma proteome.[11] The development of CE-specific data-independent acquisition (DIA)^[6c, 20]^ and Eco data-dependent acquisition (DDA)^[20a]^ recently supported with artificial intelligence (Eco-AI)^[20a, 21]^, have begun enhancing coverage in CE-ESI-MS-based single-cell workflows.^[6c, 20a, 21-22]^ Together, these advances represent previously inaccessible capabilities for probing the molecular architecture of individual cells, ushering in a new era of subcellular systems biology.

This study was initiated with dual objectives: to advance the sensitivity of subcellular MS proteomics and to apply this capability to interrogate the proteomic landscape of asymmetric cell division. This report follows the original disclosure of the technology and subcellular work in 2023.[22] Achieving these goals required overcoming the inherent sensitivity limitations of microprobe CE-ESI-MS to reliably detect within confined subcellular niches at least 1,000 proteins to mark a technical milestone. While recent developments such as Eco-DIA extended proteome coverage on CE-ESI orbitrap platforms^[6c]^, challenges remain in deciphering complex chimeric MS^2^ spectra resulting from isobaric co-isolation. To address this, we integrated CE-ESI with 3 core technical improvements enabled by timsTOF MS[23]: We adapted (1) rapid ion mobility separation to reduce spectral interference, (2) ion trapping to enhance ion signal accumulation, and (3) full-duty-cycle MS^2^ for more efficient and comprehensive peptide sequencing. The method was validated following international best-practice guidelines for single-cell proteomics, emphasizing reproducibility, sensitivity, and quantitative accuracy.^[2c]^ With this enhanced platform, we explored how an embryonic (frog) stem cell (blastomere) proteome is spatially reorganized during asymmetric division—specifically, how proteomic content is partitioned across opposing cellular poles that go on to establish divergent cell fates. This biological application is a benchmark and a test case for the analytical advances presented here.

## EXPERIMENTAL

### Chemicals and Materials

All the chemicals were reagent grade or higher. LC-MS grade acetic acid (AcOH), acetonitrile (ACN), formic acid (FA), and water (Optima) were purchased from Fisher Scientific (Fair Lawn, NJ). Ammonium bicarbonate (AmBic) was obtained from Avantor (Center Valley, PA). The HeLa proteome digest standard (part no. 88329) and MS-grade TPCK-treated trypsin (part no. 90057) were sourced from Thermo Fisher Scientific (Pierce, Rockford, IL). Microcapillaries for cell sampling and electrospray were fabricated from borosilicate glass tubing (0.75/1.00 mm inner/outer diameter, part no. B100-75-10; 0.50/1.00 mm, part no. B100-50-10; Sutter Instrument, Novato, CA) using a P-1000 micropipette puller (Sutter Instrument).

Capillaries for CE separations (fused silica, 40/105 µm inner/outer diameter, part no. 1068150596) were supplied by Polymicro Technologies (Phoenix, AZ). All the protein and peptide samples were processed in 0.2 mL LoBind microcentrifuge tubes (part no. 951010006, Eppendorf USA) to minimize sample loss due to adsorption.

### Solutions

The *cell aspiration buffer* contained 50 mM AmBic in water. The CE *background electrolyte* (BGE) was 1 M FA in aqueous 25% (v/v) ACN. The CE-ESI *sheath solution* contained 0.5% (v/v) AcOH in aqueous 10% (v/v) ACN. The proteome digests were resuspended in the *sample solvent* containing 0.05% (v/v) FA in aqueous 75% (v/v) ACN for CE-MS. 100% Steinberg’s solution was prepared with 17 % (w/v) sodium chloride, 0.5 % (w/v) potassium chloride, 0.5 % (w/v) calcium chloride, and 1.025 % (w/v) magnesium sulfate.

### Animal Care and Embryology

Adult *Xenopus laevis* frogs (Nasco, Fort Atkinson, WI or Xenopus1, Dexter, MI) were received and kept in a breeding colony. All the protocols regarding the humane care, handling, and use of *X. laevis* were approved by the Institutional Animal Care and Use Committee of the University of Maryland, College Park (approval nos. R-FEB-21-07 and R-FEB-24-05). Embryos were obtained through natural mating of multiple parental pairs to incorporate biological variability into proteomic analyses and through *fertilization in vitro* to precisely control developmental timing for the ultraviolet (UV) ventralization experiments. Two-cell embryos exhibiting stereotypical pigmentation patterns were cultured following established protocols.[24] Blastomere identities were determined based on their position, morphology, and pigmentation and assigned according to established fate maps of 8- and 16-cell stage embryos in this model system.[24–25]

### Subcellular Sampling

The cellular contents were collected *in vivo* or *in situ* via capillary microsampling, as described in our previous work.^[6a, 6b]^ Briefly, borosilicate glass capillaries were pulled to produce micropipettes with ∼20 µm diameter tips. Each micropipette was mounted on a 3-axis manual translation stage and guided under a stereomicroscope to pierce the targeted blastomeres in sufficient finesse to reproducibly sample the polar contents of the giant D1 precursor cell. A controlled vacuum pulse (–40 psi for 10 s) was applied to aspirate ∼10 nL of the cytoplasm, as confirmed via imaging of the collected cytoplasm forming a droplet in mineral oil. This sampling volume correspond to ∼5% of the cell’s cytoplasmic content (∼180 nL). Different, clean micropipettes were used to sample the future left D11 (‘D11) and future D12 (‘D12) poles in the left dorsal-animal (D1) blastomere of the 8-cell stage (n = 4 BRs) as well as the left D11 and left D12 blastomeres at the 16-cell stage (n = 4–5 BRs; **Fig. 1**). The aspirates were immediately expelled into 5 µL of 50 mM AmBic in 0.2 mL LoBind microcentrifuge tubes and processed for bottom-up proteomics.

**Figure 1.**
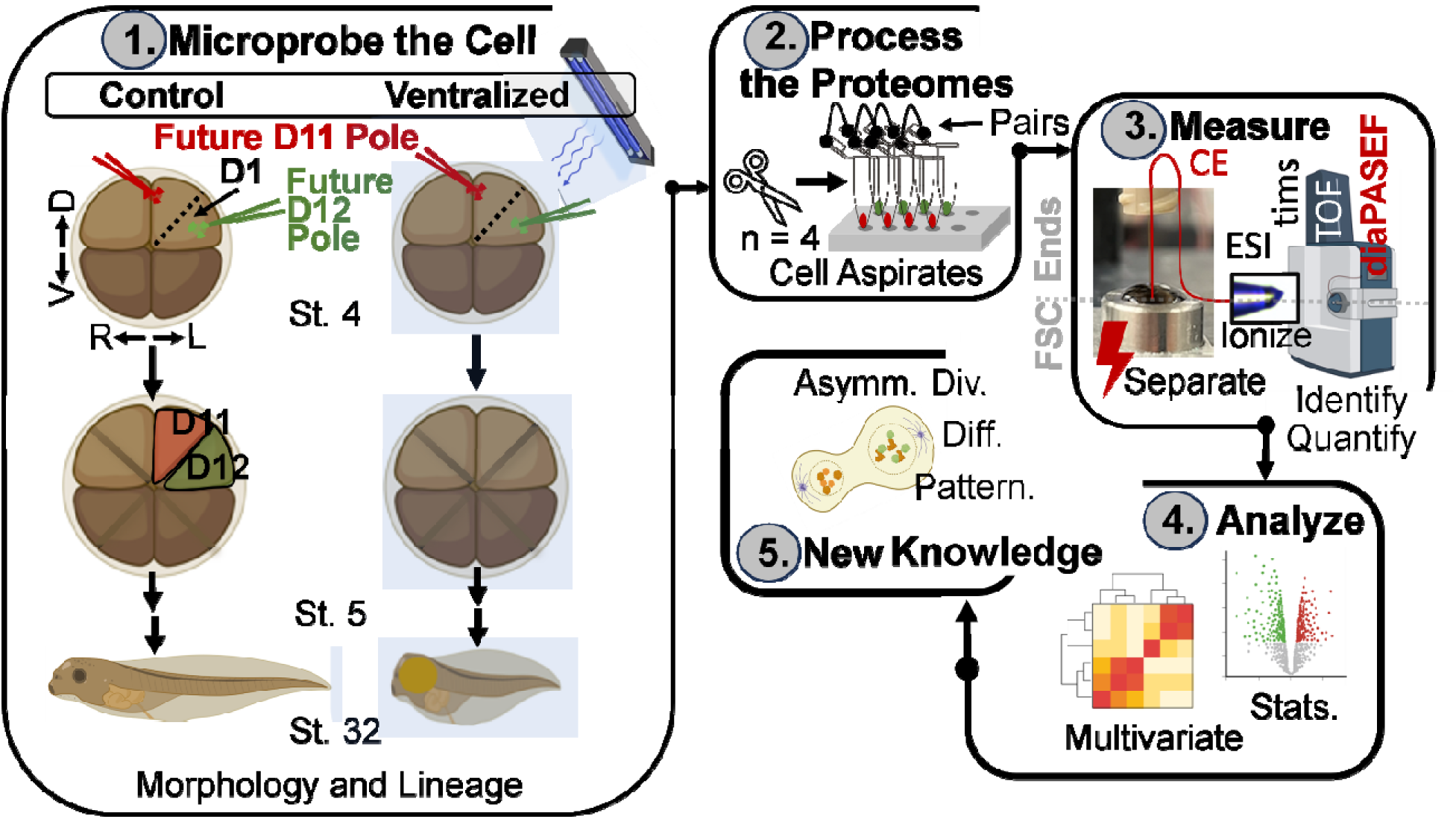
Our experimental approach to profiling asymmetric cell division in live embryos of the *Xenopus laevis* frog. Ca. 5% of the total cell volume was capillary micro-aspirated from the opposing future dorsal-animal-midline (D11) and animal-midline-lateral (D12) poles of the animal D1 cell in the live 8-cell *X. laevis* embryo, destined to form the respective neural and epidermal tissues in the larva. The contents of the descendent D11 and D12 blastomeres inheriting these polar niches were collected to trace differentiation. The proteomes were trypsin-digested through a simplified bottom-up proteomic workflow and measured on a trapped ion mobility spectrometry time-of-flight (timsTOF) mass spectrometer (timsTOF PRO, Bruker, **Methods**). CE-diaPASEF was configured for sensitivity using subcellular, ∼200 pg of the HeLa proteome digest. The blastomere proteome profiles were analyzed through multivariate and statistical means. The inheritance of the observed proteome differences was tested using ultraviolet (UV) light ventralization. The CE-diaPASEF subcellular and cellular analyses revealed complex proteome remodeling among the polar opposites establishing different fates during asymmetric cell division in the chordate embryo. (Created with BioRender.com).

### UV Ventralization

The embryos were ventralized through ultraviolet (UV) radiation exposure as per a previous protocol.[26] At 5 mins post-fertilization (pf), the embryos were dejellied and promptly transferred into Corning vials containing 100% Steinberg’s solution for UV treatment. They were then irradiated with 254 nm UV light (4 W power, part no. 95-0021-12, Analytik Jena, Upland, CA) for varying durations to optimize ventralization and survival: 0, 30, 60, 90, or 180 s. After the treatment, the embryos were either cultured to the 8-cell stage for proteomic analysis or allowed to develop to the larval stage for morphological assessment. Dorsal axial development was evaluated using the Dorsal Anterior Index (DAI), as previously described[27]. In the control group (n = 15), all embryos developed normally with a DAI of 5. Embryos exposed to 30 s of UV (n = 18) showed a mean DAI of 4.1 ± 1.0; those exposed for 60 s (n = 18) had a DAI of 1.2 ± 1.2; and those irradiated for 90 s (n = 15) yielded a DAI of 1.5 ± 0.7. All the embryos exposed to up to 90 s of UV survived to the larval stage (100% survival rate). However, only 67% of embryos irradiated for 180 s (n = 15) reached the gastrula stage. Therefore, a 90-s UV exposure was optimal on our experimental setup for efficient ventralization (DAI ≤ 2) while preserving high viability (∼100%), making these conditions suitable for downstream functional experiments.

### Proteome Preparation

The conventional steps of the bottom-up proteomics workflow, such as alkylation and reduction, were eliminated to minimize sample loss as per our validated approach.^[6b]^ The subcellular D1 proteomes were denatured by rapid heating to 60 °C for 20 min, followed by digestion with trypsin (0.5 µL of 1 µg/µL) at 37 °C over 5 h. The D11 and D12 cell proteomes were processed similarly using additional trypsin (0.75 µL of 1 µg/µL) with a longer duration (12 h). The proteome digests were vacuum-dried in the LoBind microvials at 60 °C, then stored at –80 °C until measurement. Both the subcellular and single-cell proteome digest were resuspended into the *sample solvent* to make an 0.5 µg/µL sample solution for the CE-MS analysis, each totaling ∼3.5 µL.

### CE-nanoESI-HRMS

The peptides were electrophoresed in a 1-m-long capillary at 22 kV. The separation was performed on custom-built capillary electrophoresis (CE) platform, validated elsewhere.^[6a, 11]^ The peptides were ionized using an electrokinetically pumped CE electrospray ionization interface[28] (spray potential, ∼1,000 V), operating in the cone-jet regime to maximize ionization efficiency.^[5a, 6b, 29]^ The resulting ions were analyzed on a timsTOF PRO mass spectrometer (Bruker Daltonics) executing parallel accumulation–serial fragmentation (PASEF) MS^2^ acquisition controlled by either DDA PASEF (ddaPASEF) or DIA PASEF (diaPASEF).

Experimental parameters were as follows: end plate voltage, 0 V; MS ion transfer capillary voltage, 0 V; survey (MS^1^) m/z range, 100–1,700; 1/K_0_ range, 0.6–1.6 V s cm^−2^; ramp time, 100 ms; accumulation time, 100 ms. Only the +2 and +3 ions were selected for sequencing with the following MS^2^ settings: m/z range, 100–1,700; collision energy, 20–59 eV (linear); 1/K_0_ range, 0.6–1.6 V s cm^−2^. The <u>ddaPASEF method</u> used: PASEF scans, 12; total cycle time, 1.38 s; charge, 0–5; MS^2^ threshold, 1,500 counts; target intensity, 20,000 counts. The tested <u>diaPASEF methods</u> employed the following combinations of m/z scan range, m/z isolation window (Th), and cycle time (s): 400–1,600, 50, 2.85; 400–1,200, 25, 2.74; 500–1,200, 20, 2.74; or 400–1,200, 15, 4.80.

### Data Analysis

The MS data were processed both with and without a spectral library using DIA-NN 1.8.[30] The HeLa spectral library, containing 3,759 protein groups, was generated from 28 independent analyses of 10 ng of proteome using ddaPASEF in Spectronaut 15 with the Pulsar search engine (Biognosys, Switzerland). The HeLa protein identifications were searched against the HeLa proteome without RT prediction in the control (library-free) approach (UP000005640, downloaded from UniProt in March 2022, containing 20,380 entries). For the *X. laevis* blastomeres, a hybrid spectral library was generated from 11 diaPASEF and 22 ddaPASEF files, which included 3,918 protein groups. The *Xenopus* protein identifications were searched against the *X. laevis* proteome (UP000694892, downloaded from UniProt in June 2022, containing 43,236 entries). As the Control, the library-free searches used the following settings: minimal peptide length, 5; maximal peptide length, 35; maximum missed cleavage, 2; and precursor charge, +2–4; static modification, cysteine carbamidomethylation; dynamic modifications, methionine oxidation. All the other parameters were set to default. Protein identification was filtered to <1 % false discovery rate (FDR) against the reversed-sequence decoy database for both the library-based and library-free analyses. Proteins quantified in at least 1 TR or BR are listed in this report.

### Scientific Rigor

The HeLa proteome digest was tested in n = 3 technical replicates (TRs) to develop the method. Each *X. laevis* cellular proteome sample was measured in n = 4–5 BRs in a randomized order, with each analyzed in technical duplets to triplicates. Protein identifications were filtered to <1% FDR. The common contaminant proteins were annotated and removed.

Peptide quantification was enhanced by acquiring at least 6 data points for each m/z-selected electropherogram serving as the basis of quantification. The protein concentrations were approximated using validated methods using DIA-NN. The data were normalized to the sum in the D1 subcellular samples, where local proteome amounts may be similar but vastly different in concentration profiles between the poles, and median-scaled in the D11 and D12 blastomere aspirates, where whole-cell analysis averages subcellular differences to expect similar overall concentration means. The nonparametric Mann-Whitney U test was performed in R for non-normally distributed data, with *p* < 0.05 selected to mark statistical significance.

### Safety

Standard safety protocols were observed when handling chemicals and biological samples, including capillaries and ESI emitters posing a puncture hazard. All electrically conductive components of the CE-ESI platform were grounded or shielded behind safety interlocks to prevent potential electroshocks.

### Data Availability

All MS primary files and MS^2^ spectral libraries and the HeLa and *Xenopus* proteomes were deposited in the Proteome Exchange Consortium via the PRIDE partner repository with the data set identifier PXD063566.

## RESULTS AND DISCUSSIONS

Our goal to profile subcellular proteome reorganization during asymmetric cell division was contingent on high subcellular MS sensitivity. **Figure 1** illustrates the alignment of analytical technology goals to enable subcellular proteomics in live 8-cell and single-cell analysis in 16-cell *X. laevis* embryos during asymmetric cell differentiation. As a demonstration, we set out to enhance microprobe CE-ESI^[6a, 6b]^ sensitivity to profile more than 1,000 proteins in reproducibly identifiable subcellular niches in a live blastomere in *X. laevis* embryos. We selected the left dorsal-animal (D1) cell in the 8-cell embryo as the model. The opposing subcellular poles of the D1 blastomere are inherited asymmetrically as the embryo develops to the 16-cell stage. The dorsal-animal midline (D11) offspring gives rise to neural tissues, and the dorsal-animal lateral (D12) offspring is a reproducible precursor to the head epidermis.[24] We employed multivariate data analysis to examine these proteome patterns indicative of the subcellular and cellular identities. Although the D1, D11, and D12 lineage forms on the dorsal hemisphere of the embryo, the D1’s future D11 pole and its D11 offspring are more dorsally located than the D1’s future D12 pole and D12 cell. We performed UV-ventralization studies to assess a putative connection between the observed subcellular-to-cellular protein differences from dorsal-ventral axis patterning just one-to-two cell divisions earlier. While this study is not designed to address the biological significance of the protein gradients, the data obtained here can be used to generate hypotheses and design follow-up functional experiments to test them. All protein identifications and quantification are provided as **Supplementary Information (SI)** documents accompanying this work to enhance transparency and encourage data reuse.

### Enhancement of Proteome Sensitivity

While capillary microsampling was theoretically scalable to meet our biological objectives,^[6a, 6b]^ proteome coverage remained the key limiting factor. To evaluate technical performance, we analyzed approximately 200 pg of HeLa proteome digest, ∼80% of the total protein content of a single HeLa cell (∼250 pg/cell). This digest was sourced from a reliable commercial supplier (Methods), diluted, and analyzed on our custom-built microanalytical CE platform^[6a]^, coupled to an electrokinetically pumped electrospray ionization (ESI) source operating in the cone-jet regime for optimal ionization efficiency.^[5a, 6b, 29]^ The ions were detected on an early-generation timsTOF mass spectrometer (timsTOF PRO, Bruker, **Methods**) executing ddaPASEF or diaPASEF, the methods synchronizing ion accumulation and fragmentation for 100% ion utilization.[23] This platform was chosen because it integrated rapid ion mobility separation to reduce spectral interferences, ion trapping to boost ion signal abundance, and full-duty-cycle MS^2^ PASEF to support the identification of ∼962 proteins from ∼200 pg using Eco-IMS (ddaPASEF) on the same instrument.^[20a]^

We configured the experimental parameter space affecting spectral acquisition to improve data depth and quality further. Specifically, we investigated the effects of varying the range and width of m/z isolation windows, while managing cycle time between ∼4.8–2.74 s. We evaluated the following 4 specific windowing strategies, termed as: “broad” (400–1,200 m/z range, 15 m/z window, 4.80 s cycle time), “wide” (500–1,200, 20, 2.74 s), “extra-wide” (400–1,200, 25, 2.74 s), and “super-wide” (400–1,600, 50, 2.85 s). As shown in **Figure 2A**, the average and cumulative proteome identifications across four technical replicates (each analyzing 200 pg of HeLa digest) found the “extra-wide” configuration to yield the deepest proteome coverage. Although library-free DIA processing was convenient for limited sample amounts, library-based analysis demonstrated improved depth in recent reports.[31] To test this, we generated a project-specific spectral library from 28 ddaPASEF analysis of 10 ng HeLa proteome digest in each (equivalent to ∼40 cells), which led to 1,035 proteins over 20-min CE separations.

**Figure 2.**
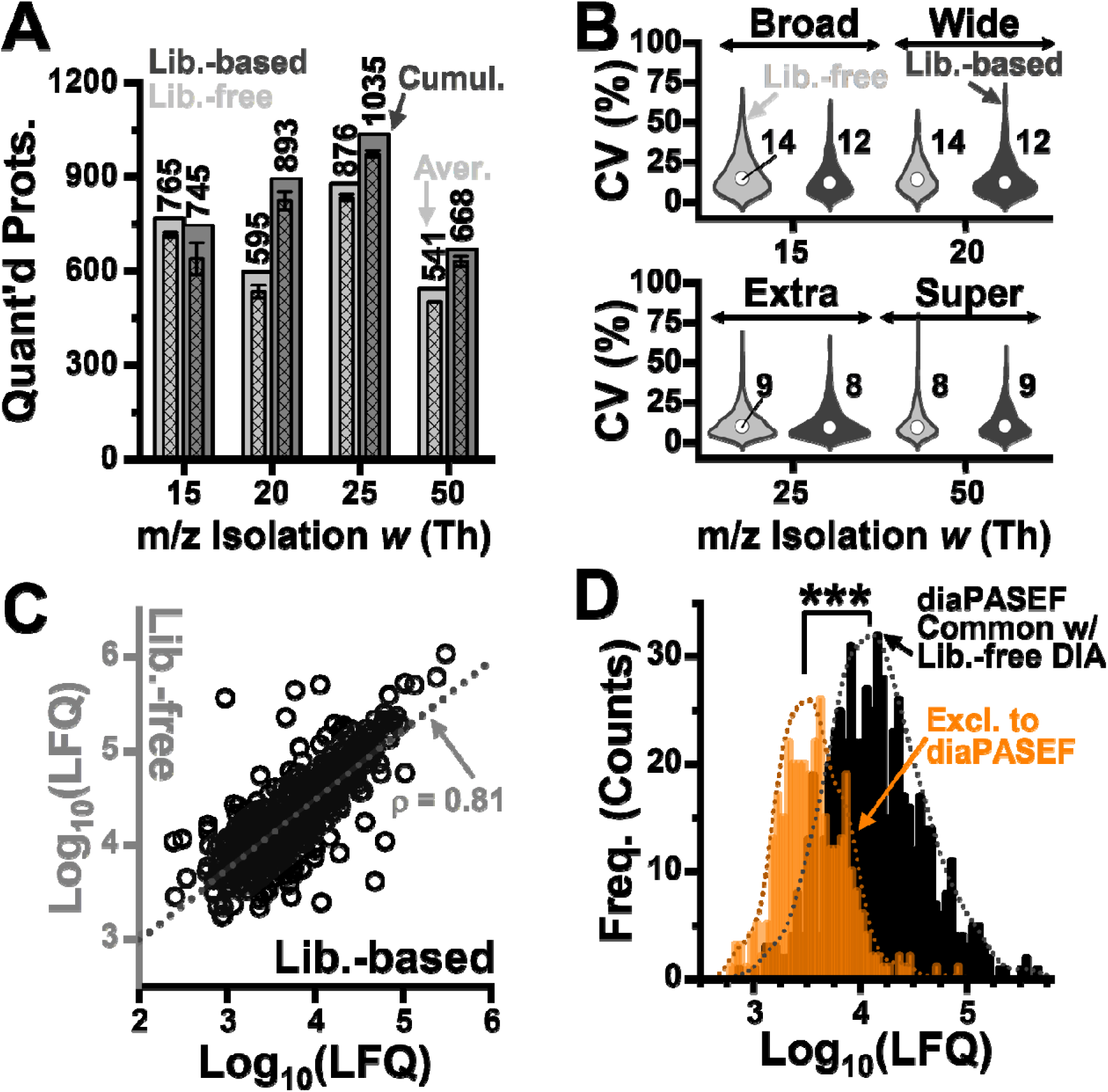
Configuration of detection sensitivity. 200 pg of the HeLa proteome digest was analyzed in technical triplicates using CE-diaPASEF, approximating to less than a cell. Different windowing strategies were tested considering m/z isolation and cycle time: broad–slow (15 Th, 4.80 s) vs fast (2.74–2.85 s cycle times) isolations across wide (20 Th), extra wide (25 Th), and super wide (50 Th) m/z ranges. **(A)** Comparison of the proteome coverage using the library-based and -free approaches. **(B)** Analysis of coefficient of variation (CV) among the DIA-NN-calculated label-free quantification (LFQ) intensities approximating concentrations, revealing robustness, with errors < 15% using both data-processing approaches. **(C)** Pearson cross-correlation analysis of protein LFQ concentration from the library-based vs -free data analyses. The data show “extra-wide” precursor ion isolation. **(D)** Comparison of protein sensitivity revealing proteins only quantifiable by the library-free diaPASEF originated from the lower regime of the measured concentration range than those mutually measured with the library-free DIA reference, confirming sensitivity enhancement. Key: ***, *p* < 0.001, Mann-Whitney.

### The Fidelity of Quantification

**Figure 2** establishes the quantifiable proteome coverage (number of quantified proteins), reproducibility (error in quantification), robustness (correlations among different studies), and sensitivity of quantification (systematic differences in concentrations). As earlier, the label-free quantification (LFQ) values were sum-normalized, log_10_-transformed, and autoscaled to estimate protein abundances on relative terms. Technical details are in the **Methods**. 1,015 proteins were quantified among the technical triplicates. These proteins are listed in **Table 1** in the **SI** (**Table S1**). **Figure 2B** depicts the coefficient of variation (CV) among the quantified protein concentrations across the technical triplicates. The median CVs were below 15% across all empirical schemes, regardless of whether library-free or library**-** based data processing was used—highlighting the method’s robustness.

We leveraged Pearson correlation analysis to quantify robustness. The proteome concentrations were implied from the quantified proteotypic peptides, using both the library-based and -free data analysis strategies. A high, ρ = 0.81, correlation coefficient revealed consistency, agreeing with our recent findings^[20b]^. In comparison to our recent CE-DIA on the a Q-Exactive plus mass spectrometer,^[20b]^ CE-diaPASEF on the timsTOF PRO identified ca. 50% more proteins from single-cell-equivalent HeLa proteome digests (∼200 pg). These newly quantified proteins populated the lower end of the dynamic concentration range (**Fig. 2D**). The improvement was statistically significant (*p* = 9.27 × 10). Based on these findings, we selected the m/z 25 diaPASEF configuration with library-based analysis for all subsequent single-cell and subcellular proteome measurements, striking a balance between depth of proteome coverage and repeatability of its quantitative profiling.

### Exploring Subcellular Proteome Niches

CE-ESI-MS was now ready for integration with capillary microsampling to explore subcellular proteome heterogeneity. As illustrated in **Figure 2**, using a stereomicroscope and manual micromanipulation, we aspirated ∼10 nL, or ∼5% of the total cell volume from the future D11 and D12 poles within the same D1 precursor blastomere of live 8-cell *X. laevis* embryos.^[6a]^ Each aspiration took under 10 s and was performed using microfabricated glass pipettes under in vivo conditions. To trace proteome inheritance, we also microsampled the descendant D11 and D12 cells *in situ,* which respectfully inherit these polar proteomes during asymmetric division from the 8-cell to the 16-cell stage embryo. Sampling was conducted in 4 embryos per condition. The identity of the cell and the analytical replicate were traced through the study. However, access to this information was blinded during unsupervised analysis and only revealed for supervised data analyses and to aid results interpretation.

The subcellular proteome samples were processed in ∼10–100-fold smaller amounts than typical in bottom-up proteomics, producing ∼3.5 µL of proteome digest per aspirate. Approximately 10 nL of each proteome extract, containing ∼0.015% of the total D1 cell proteome, was analyzed on our custom-built CE-ESI platform. In 25-min CE-MS runs using diaPASEF, 808 proteins were quantified in the future D11 pole vs. 1,022 in the future D12 pole, using a hybrid spectral library for reference (see **Table S2**). STRING network analysis revealed enrichment of pathways relating to ribosomal function, aerobic respiration, glycolysis/gluconeogenesis, proteasome activity, amino acid degradation, protein folding, and nuclear import. These processes indicate active translation and metabolic processing at the subcellular level, potentially with heterogeneity within the cell.

The proteins were quantified to measure functional activity. The protein concentrations, inferred through the calculated DIA-NN LFQ intensity, were sum-normalized, log-transformed, and auto-scaled to compare equal amounts of proteomes aspirated. Supervised statistical analysis (Student’s *t*-test, *p* < 0.05, and fold change > 1.5) revealed 158 proteins with significant differences, including 27 proteins enriched in the future D11 and 131 enriched in the future D12 pole (**Fig. 2B**, **top panel**). The remaining 755 proteins were quantified at comparable levels. The normalized protein concentrations are tabulated in **Table S3**. Gene ontology (GO) analysis using PantherDB[32] returned functions enriched in binding (86 proteins), catalytic activity (85 proteins), and structural molecule activity (20 proteins). STRING-based protein-protein interaction analysis further suggested that these proteins operate in diverse molecular contexts and subcellular localizations (data not shown).

Unsupervised hierarchical cluster analysis (HCA) of the top 45 most differentially abundant proteins corroborated the polar differences. As shown in **Figure 3B**, the TRs clustered tightly within each BR, confirming high technical reproducibility (**bottom panel**, close-up in **Fig. S1**). Importantly, the samples were grouped according to their polar identity (future D11 pole vs. future D12 pole), despite this information being withheld during data analysis. The proteins were arranged into 3 major clusters: groups #1 and #2 were enriched in the developing D12 pole, while group #3 was enriched in future D11 pole within the D1 blastomere.

**Figure 3.**
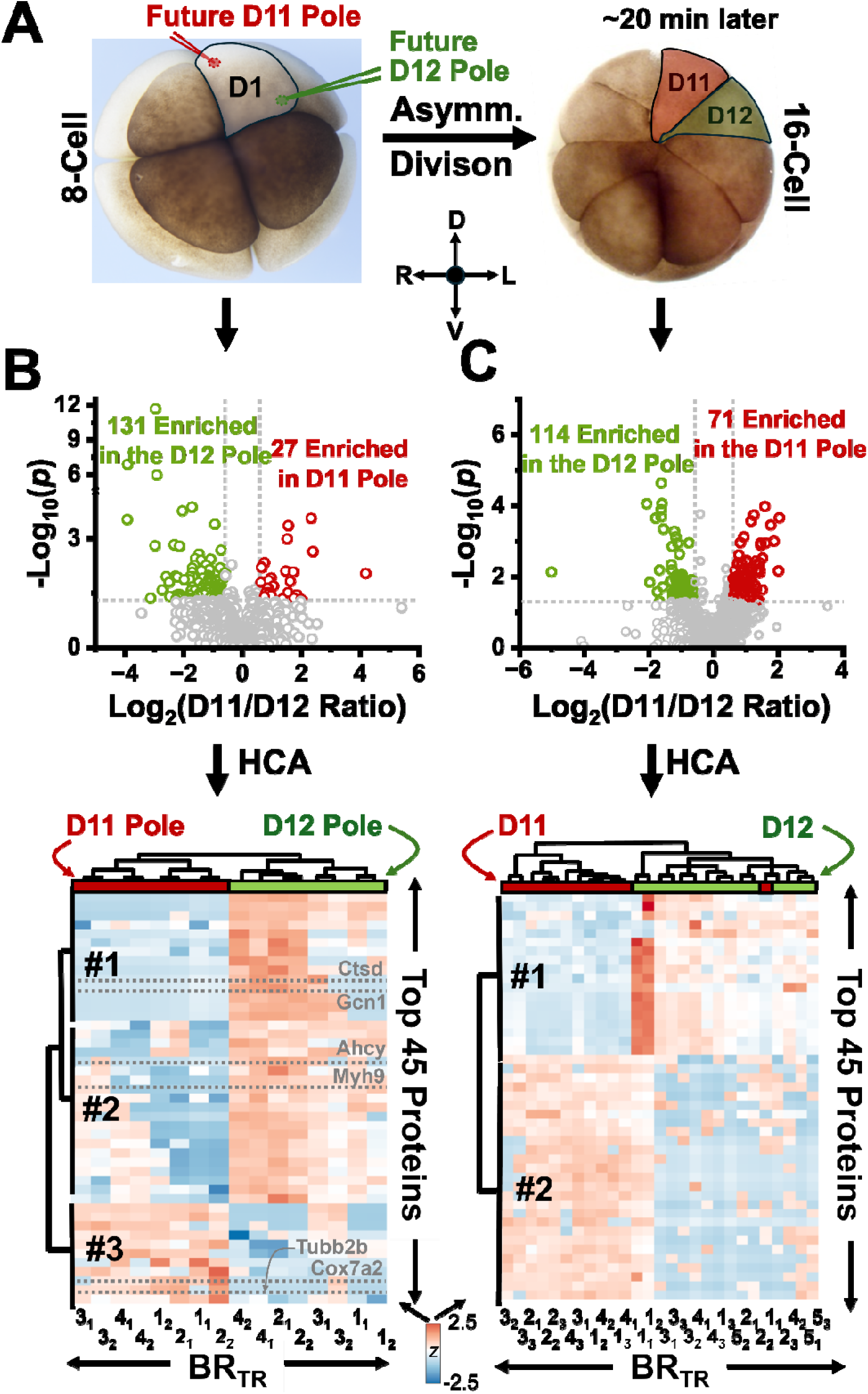
Proteome profiling of an asymmetrically dividing, identified cell in the live *X. laevis* embryo. **(A)** The future D11 and D12 poles were identified in the left dorsal-animal (D1) cell in the 8-cell embryo, and ∼5% of the cell’s volume was extracted from these opposing cellular poles. The contents of the left descendent neural-tissue-fated D11 and left head-epidermis-fated D12 cells were analyzed to profile the proteome reorganization during cell division. **(B) (Top panel)** Statistical (Student’s t-test) analysis of the DIA-NN-calculated label-free quantification (LFQ) intensities for comparing protein concentrations between the D11 and D12 poles. 755 proteins were comparable in abundance. (**Bottom panel**) Unsupervised hierarchical cluster analysis (HCA) on the normalized LFQ intensities estimating the concentrations (closeup in **Fig. S1**), revealing high technical robustness and reproducible proteome differences between the poles. **(C)** Comparison of the descendent D11 and D12 single cells’ proteome profiles using (**top panel**) the statistical (Student’s t-test) and (**bottom panel**) unsupervised HCA analyses (close-up in **Fig. S2**). Each cell sample was labeled with the identity of the embryo, biological replicate (BR), and technical replicate (TR) as follows: BR_TR_. Statistical significance is marked at *p* < 0.05 (Student’s t-test). Protein groups are labeled (#1–3 in **panel A** and #1–2 in **panel B**).

We inquired about randomly selected proteins using Xenbase.org[33]. For example, cytochrome c oxidase subunit 7A2 (Cox7a2) was 5.27-fold more abundant in the D11 pole (cluster #3; *p* = 0.002), consistent with its role in mitochondrial respiration[34] and expression in neural lineages like eye and somite during tailbud stages[35]. Class II β-tubulin (Tubb2b) showed a 5.05-fold enrichment in D11 (*p* = 0.0003), aligning with its forebrain-specific expression.[36] Conversely, myosin heavy chain 9 (Myh9) was 1.48-fold more abundant in D12 (*p* = 0.01), in line with its epidermal expression in zebrafish.[37] Additional D12-enriched proteins included cathepsin D (Ctsd, 6.02-fold, *p* = 0.0016), eIF2 alpha kinase activator homolog GCN1 (Gcn1, 4.40-fold, *p* = 0.005), and adenosylhomocysteinase (Ahcy, 2.4-fold, *p* = 0.01), all of which are linked to epidermal, retinal, or ectodermal fates,[33] a destiny of the D12 lineage[24]. These examples underscore the functional divergence of subcellular proteomes in early embryogenesis.

### Proteome Remodeling During Asymmetric Division

We hypothesized that some subcellular proteome differences observed in the D1 precursor would be retained in its descendant cells, the D11 and D12 cells, during asymmetrical division. Other proteome profiles may be altered during a dynamic interplay of molecular activation and deactivation driving cell processes. We tested the hypothesis of inheritance by extending microprobe CE-diaPASEF to D11 and D12 blastomeres in *n* = 4–5 BRs, each from a different embryo (unpaired statistical test, unlike the D1 sampling). The aspirates were randomly collected. Approximately 5 ng of total protein (∼0.03% of the cell’s proteome) was analyzed in TRs using the CE-diaPASEF method. 1,220 proteins were quantified in the D11 cell and 891 in the D12 (**Table S4**). The LFQ intensities were median-normalized, log-transformed, and auto-scaled to compare equal proteome amounts (data in **Table S5**). **Figure 2C** shows the statistical comparison of protein concentrations between the D11 and D12 blastomeres (**top panel**). While the majority (864 proteins) exhibited comparable abundance (*p* > 0.05 or fold change < 1.5), an appreciable subset, 185 proteins, were significantly differential in concentration. 114 presented higher concentration in the D12 offspring and 71 in the D11.

Unsupervised hierarchical cluster analysis (HCA) of the 45 most significant proteins revealed a clear separation between cell types. The results are shown in **Figure 3C** (**bottom panel**, close-up in **Fig. S2**). We considered the cell 2_3_ (BR_TR_) an outlier. The proteins clustered into 2 major groups: Group #1 was enriched in D12 and included yolk-related proteins such as vitellogenin a2 (Vtga2) (D12/D11 ratio = 3.09, *p* = 0.0001) and vitellogenin b1 (Vtgb1) (D12/D11 ratio = 3.47, *p* = 0.0002), consistent with the accumulation of yolk more in ventral tissues in *X. laevis*.^[5a, 5b, 38]^

In contrast, Group #2 comprised proteins enriched in D11, many associated with neural development—reflecting this lineage’s dorsal, neuroectodermal fate. Notable examples include programmed cell death protein 4 (Pdcd4, D11/D12 ratio = 2.47, *p* = 0.004), expressed during neural plate and placode formation (NF stages 15–32),[39] and developmentally regulated GTP-binding protein 1 (Drg1, D11/D12 ratio = 1.56, *p* = 0.01), expressed in the eyes, brain, and spinal cord.[40] These results highlighted complex proteome changes during cell division.

We systematically compared the proteome ratios along the dorsal-ventral direction as the analysis transitioned from subcellular poles to the whole cell. The dorsal-to-ventral protein ratios are correlated before and after division in **Figure 4A** (data in **Table S6**). Among the 829 proteins quantified across both stages, 517 did not change significantly (*p* ≥ 0.05), clustering near the origin (grey circles). In contrast, 37 proteins showed significant differences (*p* < 0.05) at both timepoints (red circles), distributed across four quadrants. Quadrant #1 contained proteins enriched in D11 both before and after division, such as Cox7a2 and G3bp1, implicated in neural tissue[35], with an ability to promote central nervous system axon regeneration[41]. Proteins in Quadrant #3 had higher levels in the D12 offspring at both stages (e.g., Ctbs), associated with endodermal and gut tissues. Quadrants #2 and #4 listed (19) proteins whose relative abundances reversed along the dorsal-ventral subcellular axis post-division, indicating dynamic remodeling. For example, G3bp2, involved in neuronal maintenance[42], was initially enriched in the D12 pole but later became more abundant in the neural-fated D11 daughter. Additionally, 152 proteins became differentially expressed only after division (yellow circles), while 123 were initially different but later equalized (blue circles). These findings support an active reorganization of spatial proteomic heterogeneity within the asymmetrically dividing cell.

**Figure 4.**
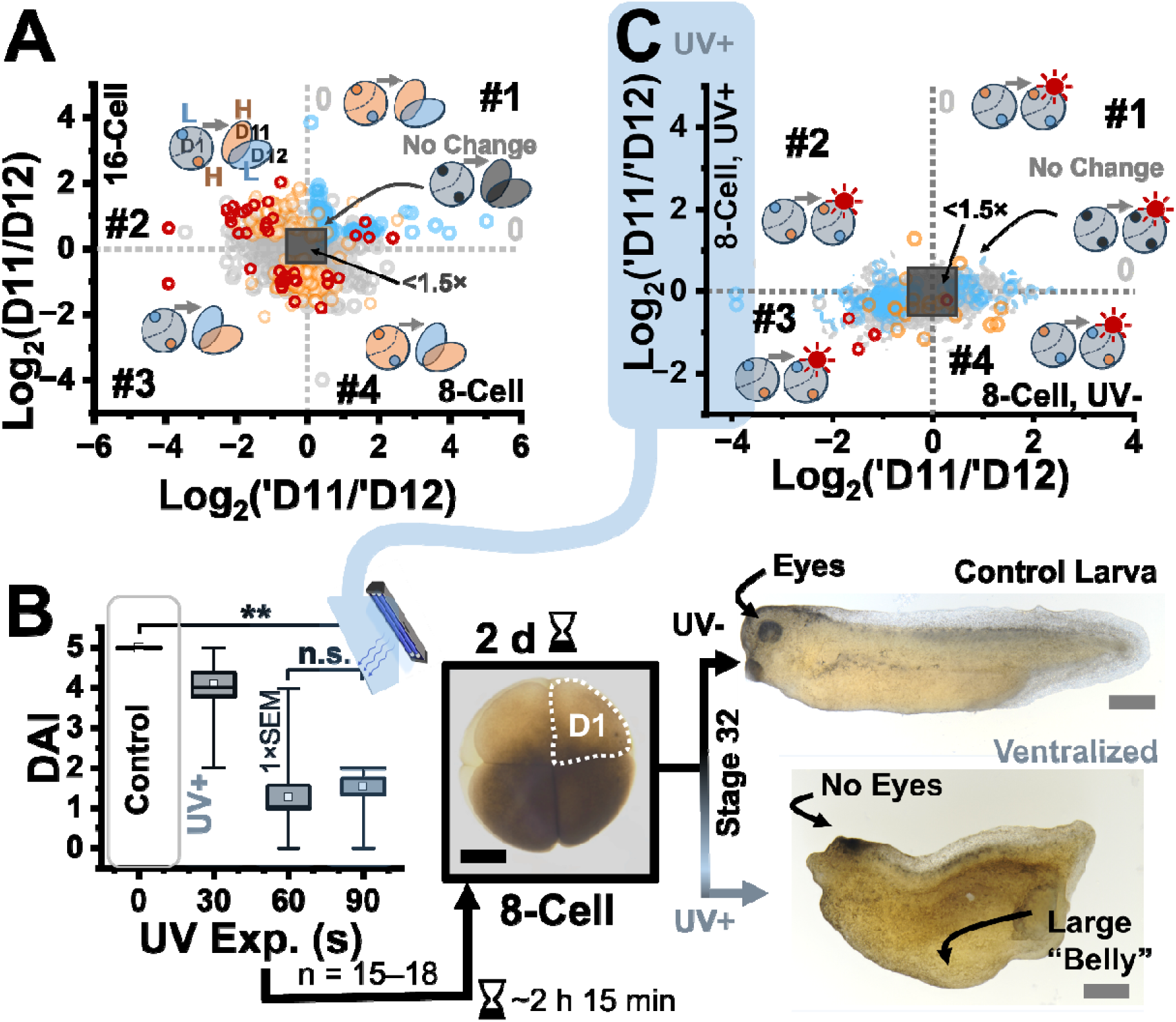
Functional testing of dorsal-ventral origin among the proteome profiles using ultraviolet (UV) ventralization. **(A)** The ratio of each protein was compared between the D11 vs. D12 cell in the 16-cell embryo and the future D11 pole (‘D11) and future D12 pole (‘D12) in the 8-cell precursor. The quantitative data revealed proteome heterogeneity along the future D11-to-future D12 axis in the D1 precursor cell. Key to color code: orange, high abundance (H); blue, low abundance (L). Key to statistical significance (data color, Student’s t-test): red, *p* < 0.05 before and after division; yellow, p < 0.05 after division or UV+ only; blue, *p* < 0.05 before division or UV+; and grey, p ≥ 0.05. **(B)** This D11–D12 axis laterally bisected the dorsal-ventral plane of the embryo, which we functionally tested via UV ventralization. N =15–18 embryos (biological replicates) were exposed to UV light 5 min after fertilization (UV+, represented by “sun”). The resulting 8-cell embryos looked indistinguishable from the control siblings (untreated, UV–). Still, they displayed hallmarks of ventralization by Nieuwkoop-Faber stage 32, as quantified using the dorsal-anterior index (DAI: 1, complete ventralization to 5, none). Missing eyes informed of perturbed neural tissue development and growth of a large ventral structure, the stereotypical “belly piece” revealed perturbed dorsal-ventral patterning. Examples of a larva from clutch-mate controls (DAI = 5) after 90-s of UV treatment (DAI = 2). Key: ***, *p* < 0.001, Student’s t-test; SEM, standard error of the mean (1× displayed). Scale bars, 250 µm (black), 1 mm (gray). **(C)** Subcellular proteome profiling following UV-ventralization (UV+, represented by “sun”) vs. the unperturbed control (UV–) at the 8-cell stage (data in **Table S8**). Multiple proteins in the D11 pole responsive to the UV light intervention were represented in pathways involved in neural tissue development.

### Functional Test of an Asymmetric Proteome Dowry

We asked whether some of the observed proteome asymmetry was imprinted by earlier dorsal-ventral patterning. In *X. laevis*, ultraviolet (UV) light exposure disrupts the transfer of dorsalizing morphogens from the ventral pole to the dorsal, producing embryos lacking head and neural tissues but enriched in ventral structures like the characteristic “belly piece”.[27] As shown in **Figure 4B**, 15/15 embryos exposed to UV for 90 s (UV+ group) developed to stages 5–6 (∼2 h 15 min pf) that were morphologically indistinguishable from the unexposed (UV-group) siblings. By the tadpole stage (stage 32), severe ventralization was evident based on lacking eyes and brains (DAI = 1.5 ± 0.7). These results validated the robustness of UV perturbation to support our functional study (**Fig. 4B**).

The future D11 and D12 poles were analyzed after UV-ventralization in n = 4 embryos, each analyzed in technical duplicates. 875 proteins were identified in the D11 pole and 832 in the D12 niche. The identified proteins are listed in **Table S7**. As earlier, the proteins were quantified using LFQ via DIA-NN. The data were normalized to the sum, before log_10_-transformation, and auto-scaling. **Figure S3** shows the HCA of the top 50 proteins. The technical duplicates were clustered together, confirming high technical reproducibility. A clear separation emerged between the poles, indicating that subcellular proteome differences persisted after UV-ventralization.

We asked how the D11-D12 subcellular profiles responded to ventralization. As earlier, the statistical (**Fig. S4**) and unsupervised HCA analyses (**Fig. S5**) revealed high technical repeatability and differences between the control and UV ventralized embryos. **Figure 4C** correlates the relative ratio of the future D11 and future D12 pole in the 8-cell embryo before UV– group) and after UV treatment (UV+ group). Among 609 proteins quantified between the conditions, 467 were unaffected by UV exposure (*p* ≥ 0.05, grey circles). A gray square shows proteins with fold change less than ∼1.5 to select potential biological importance on the basis of <15% CV error we earlier observed in quantification (recall **Fig. 2B**). Interestingly, 118 differentially expressed proteins in the control (UV–) became indistinguishable after UV treatment (blue circles), suggesting some relationship with dorsal-ventral patterning. For example, Cox7a2, previously enriched in D11 (D11/D12 ratio = 5.27, *p* = 0.002), became equally abundant post-UV (D11/D12 ratio = 0.95, *p* = 0.617), as did Tubb2b (D11/D12 ratio reduced from 5.05 to 1.04). The relative profiling of the dorsal-ventral poles, including many known markers of neuroectodermal tissues[35, 43], are summarized in **Table S8**. STRING protein-protein interaction analysis revealed these proteins to generally associate with biosynthesis of amino acids, respiratory electron transport chain, fatty acid degradation, and protein folding.

We used the data to inspect the proteome changes on the most dorsal side of the embryo, the D11 niche, which was anticipated to be most sensitive to ventralization. We profiled the D1 cell’s proteome profile following UV-ventralization. The LFQ data from the library-based search was sum-normalized, log_10_-transformed, and auto-scaled for each BR. **Figure S6** provides the quantitative comparison of the normalized intensities. Of 927 total proteins quantified, 734 were unperturbed by light treatment. 83 proteins were upregulated, and 110 were downregulated following UV treatment in the D11 pole (**Fig. S6).** The data is available in **Table S9**.

This proteome reorganization was annotated against canonical knowledge. We employed Ingenuity Pathway Analysis (IPA) to inquire about the function of the proteins most diminished upon ventralization to the D1 blastomere’s D11 pole. The *Xenopus* proteomes were mapped to the Human orthologs for this portion of the study (**Table S9**). Broad terminologies related to cellular assembly and organization (IPA score = 55), cell signaling (score = 39), and cell morphology (score = 17) were diminished. Embryonic development and nervous system development was also detectably suppressed (score = 10). Among the downregulated proteins, 27 had corresponding mRNAs highly expressed in neural tissues such as the brain, eye, and spinal cord—regions that arise from the D11 lineage. For instance, Ranbp1, essential for neural crest migration [44], was significantly decreased (D11/D12 ratio = 0.64, *p* = 0.04), as was Nap1l1, a regulator of neural progenitor proliferation[45] (ratio = 0.61, *p* = 0.002). Notably, the 14-3-3 proteins Ywhae, Ywhaq, and Ywhaz, all key to neuronal development,[46] were also diminished (ratio = 0.38–0.60, *p* = 0.002–0.04). An interaction analysis of these proteins is presented in **Figure S7**. These proteins were in changes were potentially informative of the mechanism of dorsal-ventral patterning. While the differentially regulated proteins were not directly annotated as components of the dorsal-ventral pathways, several play roles that intersect with or influence Wnt/BMP signaling and, consequently, dorsal-ventral patterning based on the canonical knowledge. These include but are not limited to signal transduction via cytoskeletal regulation (Actb), cell signaling and membrane dynamics (Anxa1, Anxa 5), epigenetic regulation (Asf1a), energy production (multiple Atp5 synthase subunits). For example, ubiquitin C-terminal hydrolase L1 (Uchl1), which stabilizes β-catenin by inhibiting its degradation via deubiquitination,[47] was significantly downregulated (ratio = 0.29, *p* = 0.0006). Similarly, S-phase kinase-associated protein 1 (Skp1), a β-catenin stabilizer,[48] had diminished concentration. Notably, both overexpression and knockdown of the *skp1* gene have been shown to impair neural crest induction during *Xenopus* development.[48] Future studies are needed to examine the biological implications of the many protein differences uncovered in this work using subcellular CE-diaPASEF MS proteomics.

## CONCLUSIONS

This study represents a significant advance in the sensitivity and application of subcellular MS for studying the proteomic landscape of asymmetric cell division. Using as little as ∼200 pg of proteome digest, the equivalent to ∼80% of the proteome content of a single HeLa cell, we consistently identified over 1,000 proteins with high quantification reproducibility (coefficient of variation <15%), both with and without a spectral library. The CE-diaPASEF platform maintained this high sensitivity in subcellular proteome analyses from opposing poles of the D1 blastomere in the 8-cell *Xenopus laevis* embryo, which divides asymmetrically to produce the neural-tissue-fated D11 and head-epidermis-destined D12 offspring cells. Furthermore, UV-induced ventralization demonstrated that some proteomic asymmetries—including those involving the β-catenin/Wnt signaling pathway—depend on dorsal-ventral patterning, supporting their functional role in early cell fate decisions.

Emerging technologies are poised to further enhance the scope and impact of subcellular CE-diaPASEF proteomics. Innovations such as automated microsampling platforms capable of large-volume sample stacking[49] on limited proteomes (e.g., RoboCap[49]), multiplexed quantification strategies using full^[5a]^ or protein-depleted carrier proteomes[50], and next-generation mass spectrometers with ever-increasing speed and sensitivity will extend the analytical reach of this method. Subcellular patch-clamp proteomics[11], particularly for probing neuronal compartments, stands to benefit greatly from such improvements. Additionally, integration with high-sensitivity nanoLC-based workflows, such as SCoPE-MS^[10a]^, nanoPOTS[51], oil-air-droplet (OAD)[52], and whole-neuron patch-clamp proteomics[12], offers complementary approaches for deep profiling of limited samples.

As these technical capabilities evolve, they will open new frontiers for exploring proteome heterogeneity at subcellular resolution. We envision microprobe CE-diaPASEF to be scalable to smaller dimensions and cells, where precision control of microprobe sampling and (endogenous/exogenous) organ-specific optical labels can aid the targeting accuracy to specific organelles. Together, enhanced sampling, separation, and detection strategies will deepen our understanding of how protein localization shapes cellular identity and function—within the confines of the cell.

## ACKNOWLEDGMENT

This research was partially supported by the Arnold and Mabel Beckman Foundation (Beckman Young Investigator award to P.N.), the National Institute of General Medical Sciences of the National Institutes of Health (award no. R35GM124755 to P.N.), and the COSMOS Club Foundation (fellowship to B.S.). We thank M. Tarikul Islam for assistance during data analysis using IPA.

---

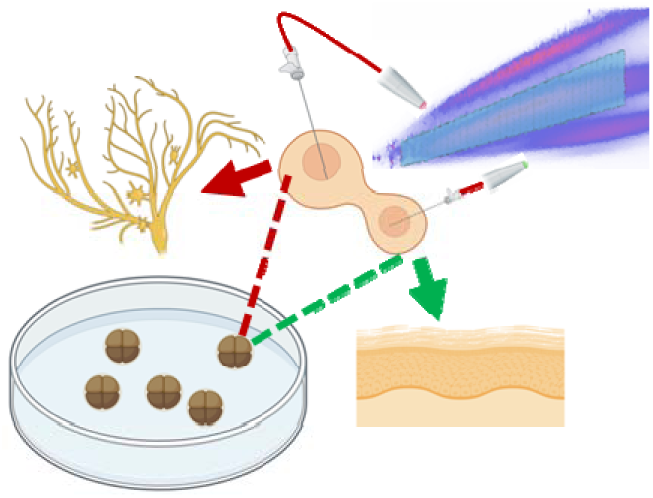
Table of Content Graphic.

## Notes

https://www.ebi.ac.uk/pride/

### Competing Interest Statement

The authors have declared no competing interest.

