## Supplementary Document for "Subcellular Mass Spectrometry Reveals Proteome Remodeling in an Asymmetrically Dividing (Frog) Embryonic Stem Cell"

### FIGURES

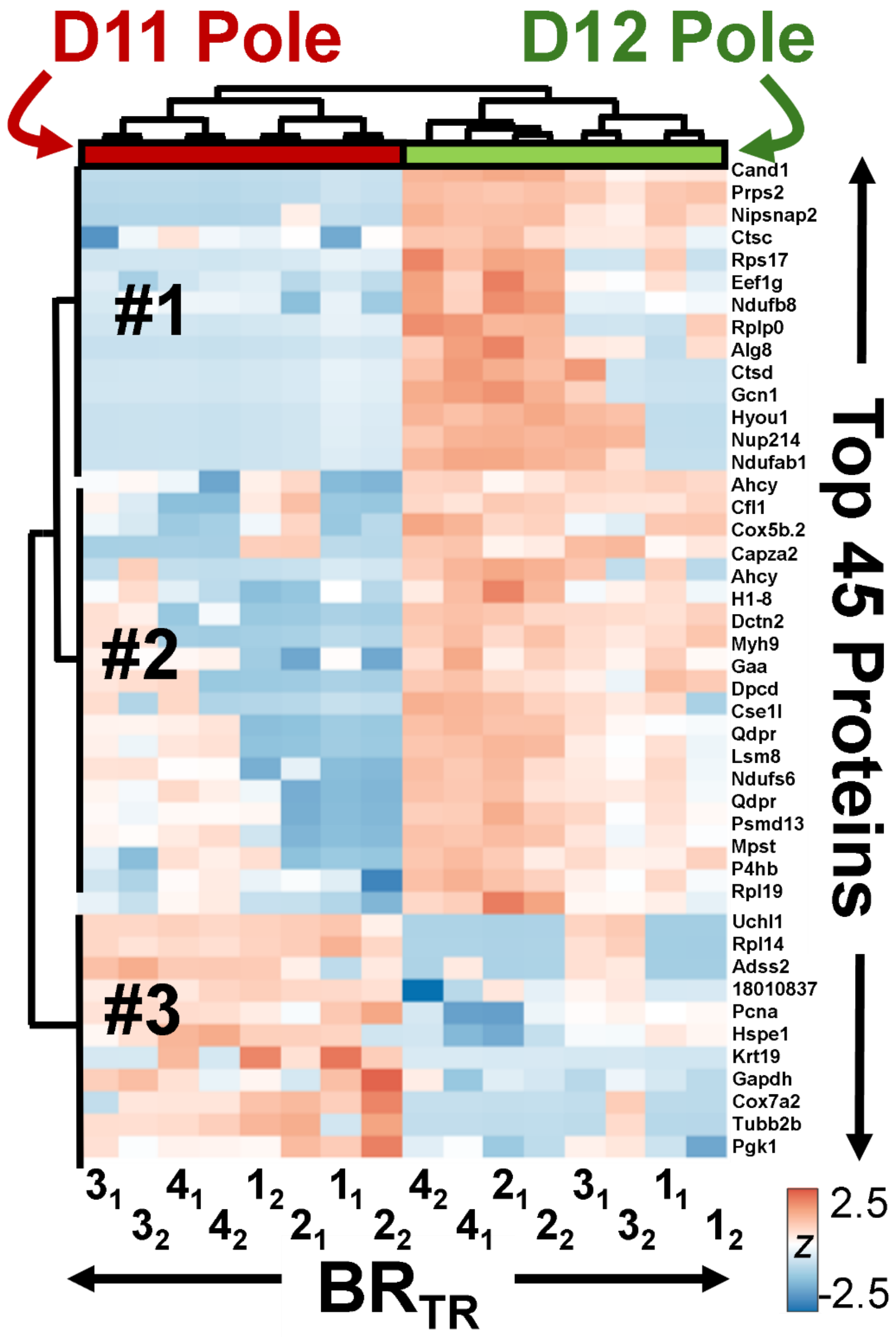
D11 Pole
D12 Pole
#1
#2
#3
Cand1
Prps2
Nipsnap2
Ctsc
Rps17
Eef1g
Ndufb8
Rplp0
Alg8
Ctsd
Gcn1
Hyou1
Nup214
Ndufab1
Ahcy
Cfl1
Cox5b.2
Capza2
Ahcy
H1-8
Dctn2
Myh9
Gaa
Dpcd
Cse1l
Qdpr
Lsm8
Ndufs6
Qdpr
Psmd13
Mpst
P4hb
Rpl19
Uchl1
Rpl14
Adss2
18010837
Pcna
Hspe1
Krt19
Gapdh
Cox7a2
Tubb2b
Pgk1
Top 45 Proteins
$3_1$ $3_2$ $4_1$ $4_2$ $1_2$ $2_1$ $1_1$ $2_2$ $4_2$ $4_1$ $2_1$ $2_2$ $3_1$ $3_2$ $1_1$ $1_2$
$BR_{TR}$
2.5
Z
-2.5

**Figure S1**. Close-up of the HCA-heat map of the top 45 most significantly differential proteins between the D11 pole and D12 pole of the D1 precursor cell in the 8-cell *X. laevis* embryos (**Fig. 3C**, **left panel**). Each protein is labeled with their corresponding gene names.

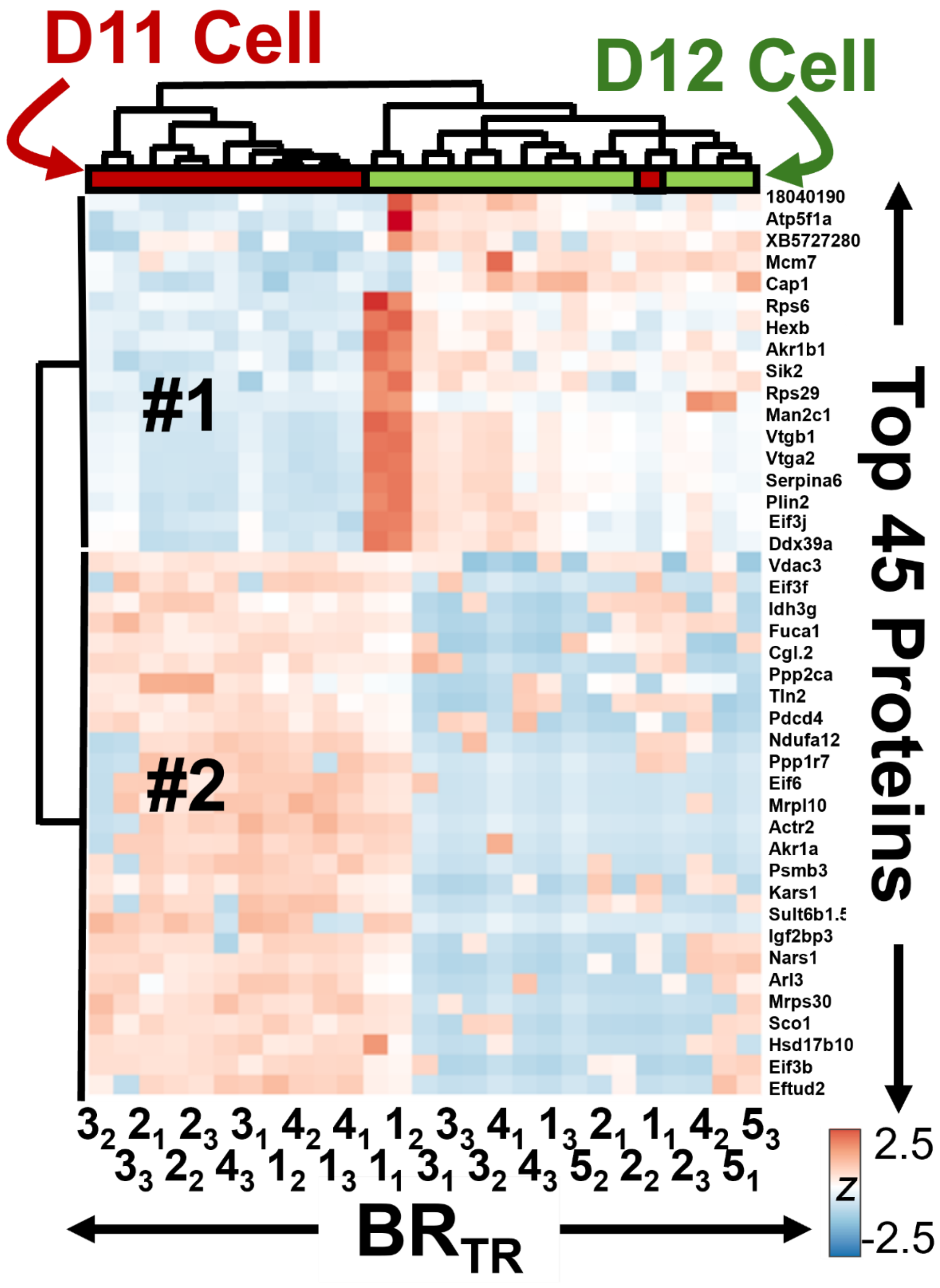

**Figure S2**. Close-up of the HCA-heat map of the top 45 most significantly differential proteins between the D11 and D12 cells in the 16-cell X. laevis embryos (**Fig. 3C, right panel**). Each protein is labeled with their gene name.

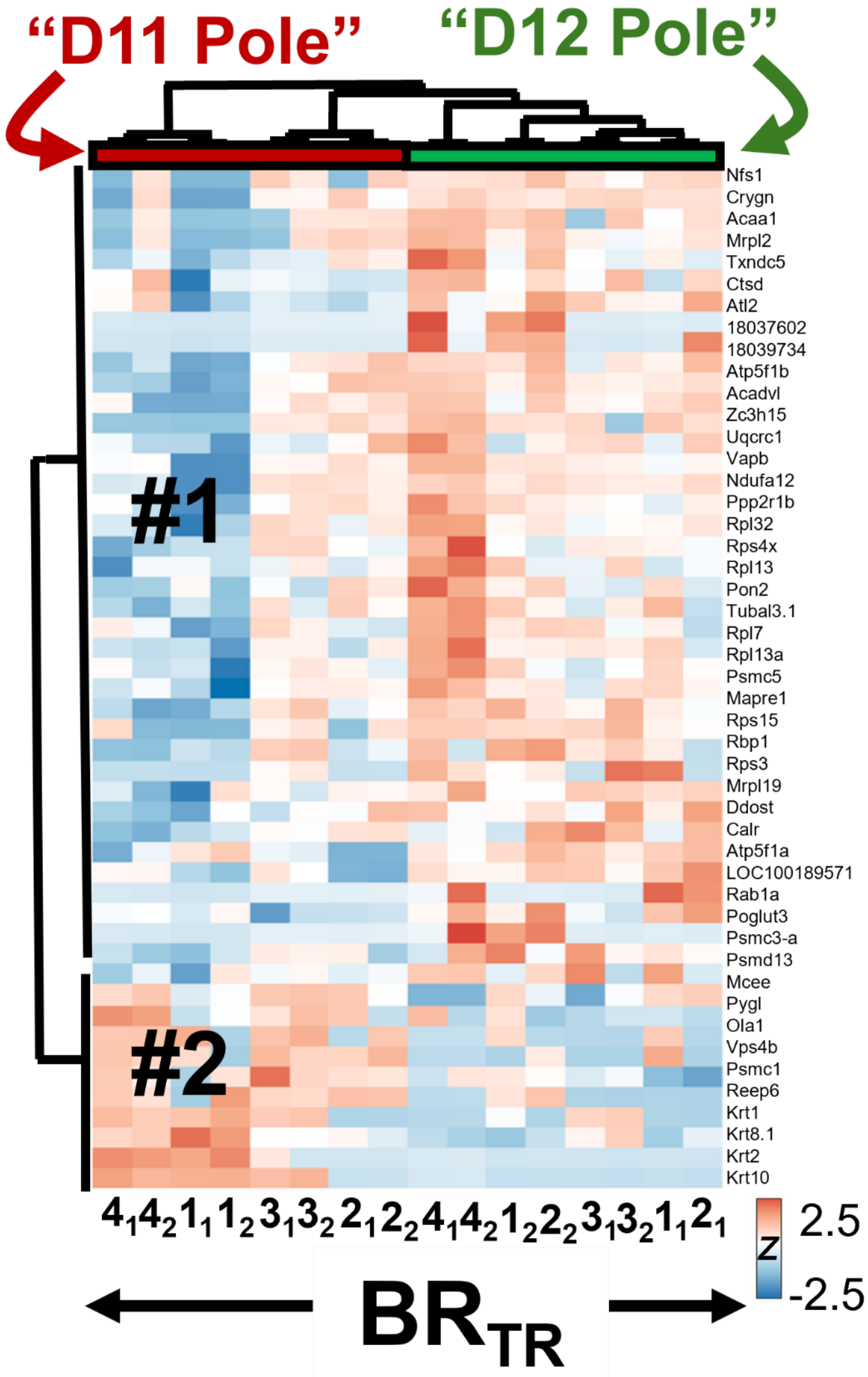

**Figure S3.** The HCA-heat map of the top 50 most significantly differential proteins between the D11 poles and D12 poles after UV ventralization at the 8-cell stage of *X. laevis* embryos.

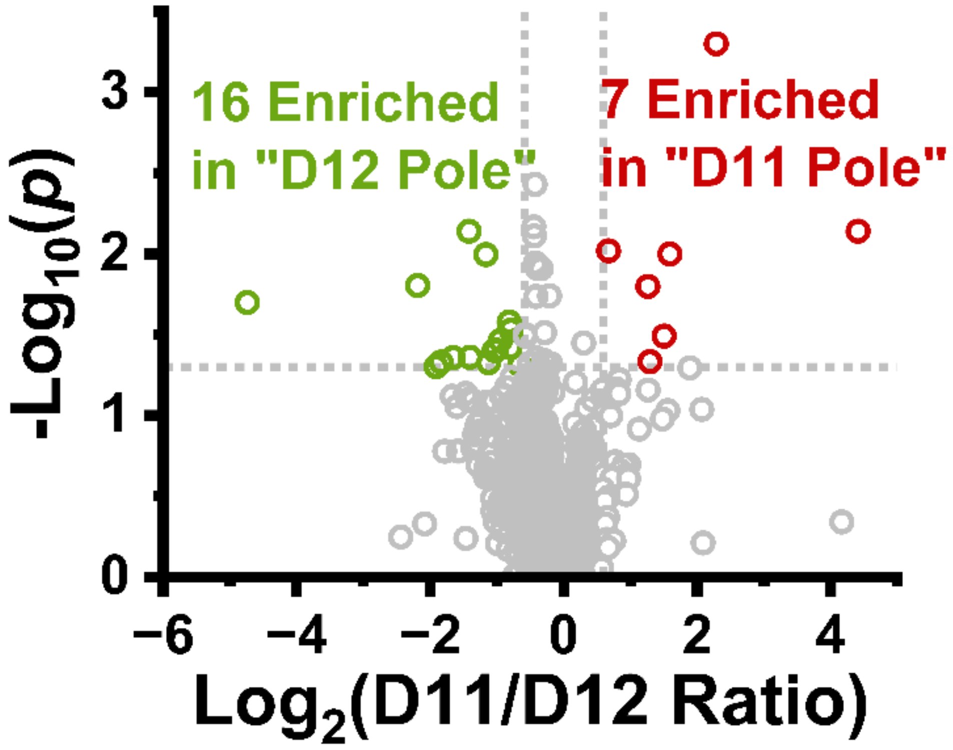

**Figure S4.** Statistical analysis of the observed protein LFQ concentrations between the "D11" and "D12" poles in the D1 precursor blastomere after UV perturbation (UV+).

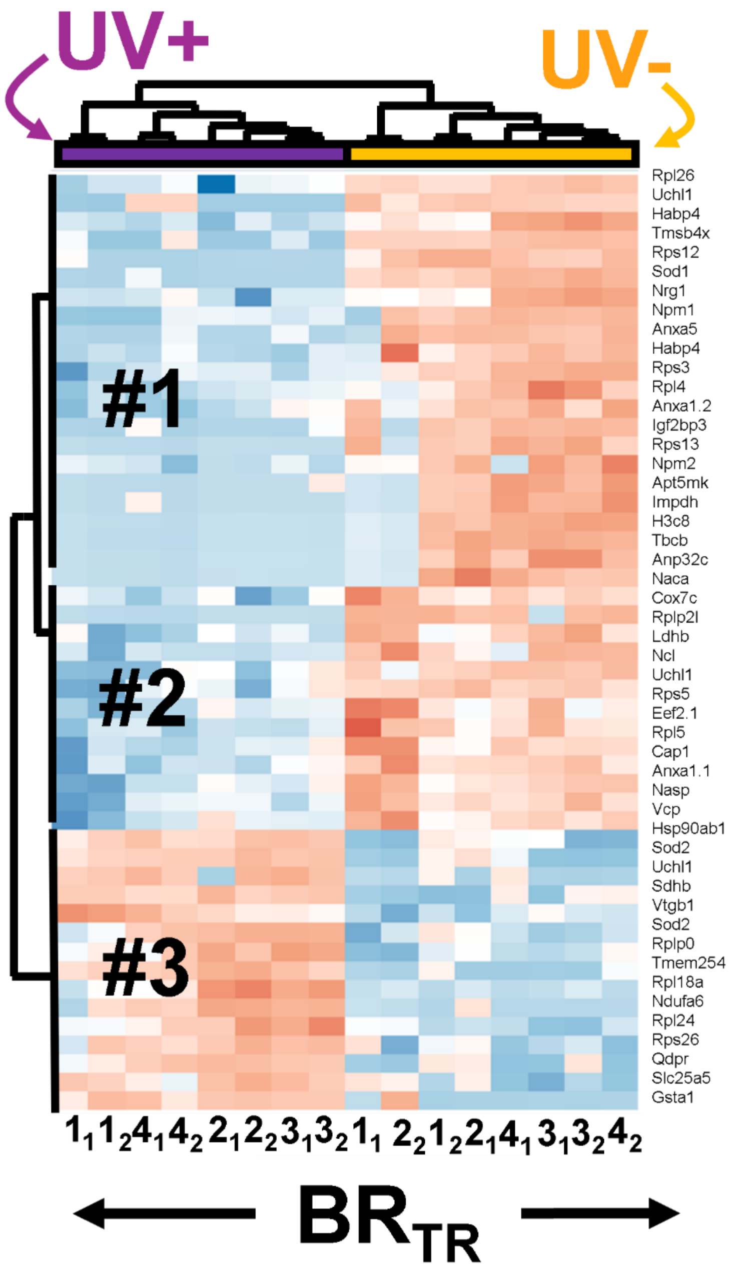

**Figure S5.** The HCA-heat map of the top 50 most significantly differential proteins between before vs after UV ventralization in the D11 poles at the 8-cell stage of *X. laevis* embryos.

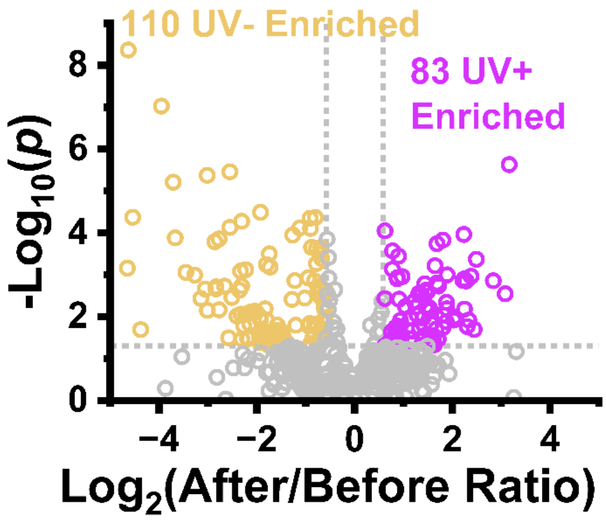

**Figure S6.** Statistical analysis of the observed protein LFQ concentrations in the D11 poles before and after UV perturbation (UV+).

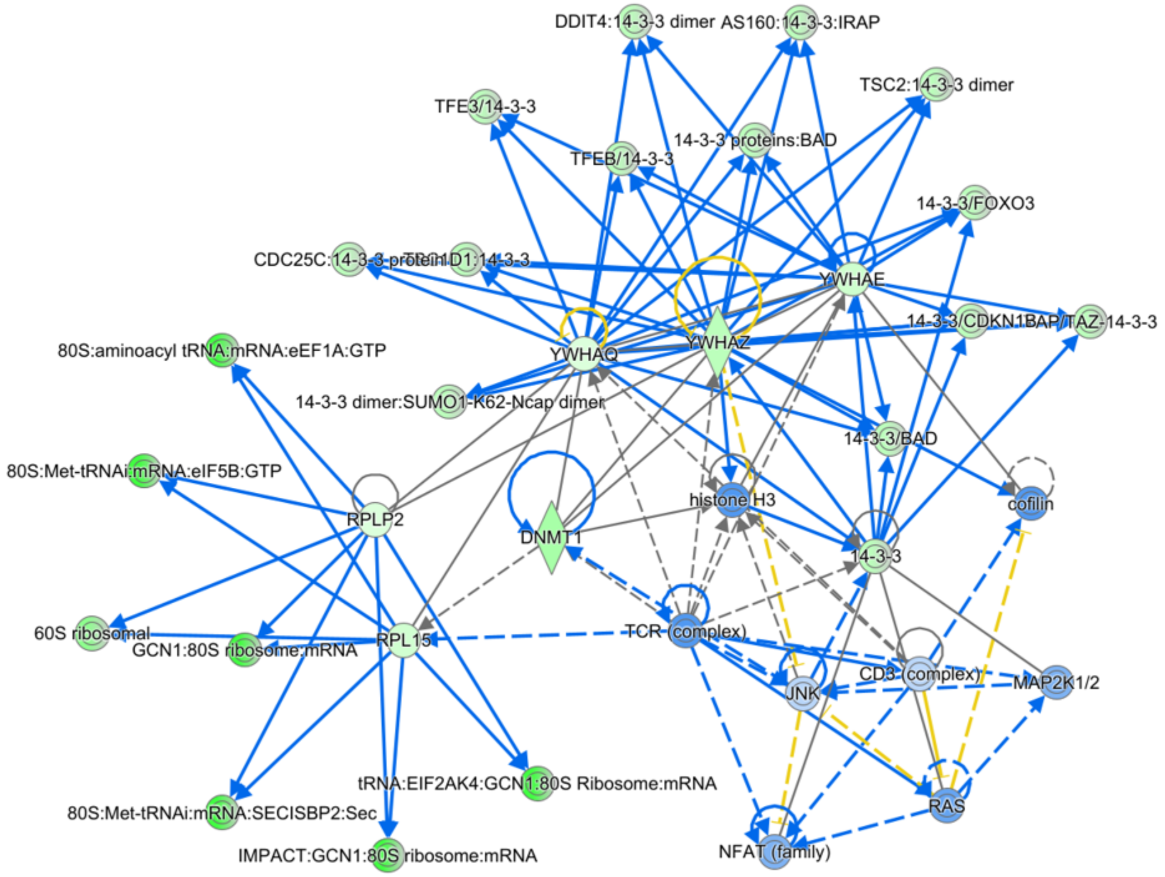

**Figure S7.** Ingenuity pathway analysis (IPA) of proteins downregulated upon ultraviolet ventralization with canonical functions in embryonic and nervous system development. Human protein ortholog genes are shown.
